# Proteome Exploration of *Legionella pneumophila* for Identifying Novel Therapeutics: A Hierarchical Subtractive Genomics and Reverse Vaccinology Approach

**DOI:** 10.1101/2020.02.03.922864

**Authors:** Md Tahsin Khan, Araf Mahmud, Mahmudul Hasan, Kazi Faizul Azim, Musammat Kulsuma Begum, Arzuba Akter, Shakhinur Islam Mondal

**Author notes:** Corresponding author Dr Md Shakhinur Islam Mondal, Associate Professor, Department of Genetic Engineering and Biotechnology, Shahjalal University of Science and Technology, Sylhet-3114, Bangladesh.

## Abstract

*Legionella pneumophila*, the causative agent of a serious type of pneumonia (lung infection) called Legionnaires’ disease. It is emerging as an antibacterial resistant strain day by day. Hence, the identification of novel drug targets and vaccine candidates is essential to fight against this pathogen. Herein attempts were taken through subtractive genomics approach on complete proteome of *L. pneumophila* to address the challenges of multidrug resistance. A total 2930 proteins from *L. pneumophila* proteome were investigated through diverse subtractive proteomics approaches, e.g., identification of human non-homologous and pathogen-specific essential proteins, druggability and ‘anti-target’ analysis, prediction of subcellular localization, human microbiome non-homology screening, protein-protein interactions studies in order to find out effective drug and vaccine targets. Only 3 were identified that fulfilled all these criteria and proposed as novel drug targets against *L. pneumophila*. Furthermore, outer membrane protein TolB was identified as potential vaccine target with better antigenicity score and allowed for further *in silico* analysis to design a unique multiepitope subunit vaccine against it. Antigenicity and transmembrane topology screening, allergenicity and toxicity assessment, population coverage analysis, and molecular docking approach were adopted to generate the most potent epitopes. The final vaccine was constructed by the combination of highly immunogenic epitopes along with suitable adjuvant and linkers. The designed vaccine construct showed higher binding interaction with different MHC molecules and human immune TLR2 receptors with minimum deformability at molecular level. The translational potency and microbial expression of the vaccine protein was also analyzed using pET28a(+) vector. The present study aids in the development of novel therapeutics and vaccine candidates for efficient treatment of the infections caused by *Legionella pneumophila*. However, further wet lab-based investigations and in vivo trials are highly recommended to experimentally validate our prediction.

## Introduction

*Legionella pneumosphila* is a human pathogen that is distributed worldwide within freshwater and biofilms. [1]. Inhalation of *Legionella spp.* cause Legionnaires’ disease (LD), a severe form of pneumonia and respiratory tract infection [2] and about 90% are caused by *L. pneumophila* species [3]. *L. pneumophila* belongs to the responsibility of maximum incidences regarding water-associated diseases in the United States [4,5] and caused approximately half of all reported waterborne disease outbreaks in 2005–2006 [no6,7]. *L. pneumophila* was declared as an important pathogen by the US Environmental Protection Agency (EPA) Candidate Contaminant List (CLL) due to the prevalence and seriousness of its effects on disease causing [8]. In case of disease progression, Icm/Dot type IVb secretion systems are used by *Legionella spp* in order to translocate a large repertoire of effector proteins inside the host cell cytosol [9-13], which modulate the fate of phagocytic vacuoles by preventing the fusion of phagosome-lysosome along with vacuole acidification, and by recruiting vesicles that delegate on it the properties of the endoplasmic reticulum [14]. Bacteria then damage, even completely destroy the host cell after the proliferation in this compartment.

The emergence of resistance against antibiotics or a combination of antibiotics of various infectious agents poses global threats in human health [15]. At present, the majority of pathogens causing infectious diseases are resistant to more than one drug [16]. Recent epidemiological research found *L. Pneumophila* strains display a high prevalence of tolerance (50% -100%) to widely used antibiotics comprising azithromycin, ceftriaxone, rifampicin, tigecycline, ciprofloxacin, moxifloxacin, doxycycline, erythromycin, levofloxacin and clarithromycin[17,18]. This emphasizes the urgency of developing new therapeutic antibacterial agents directing towards novel targets [15,19]. Therefore, it is exigent to identify novel therapeutic targets of *L. pneumophila.*

Identification of novel drug targets and vaccine candidates leads to drug and therapeutics component designing. The disease-based approach of the traditional method of drug discovery is both expensive, and time-consuming since significant time and devoted researchers are required to identify potential ligands. In recent years, computational aided drug targets discovery methods reduce consumption of time by eliminating compounds in this process that have a limited chance of success [20,21,22,23]. Among various computational methods and strategies, subtractive genomic analysis can be regarded worthy since data regarding genome and proteome are available in various online based databases. Subtractive and comparative microbial genomics are being used for the identification of targets in a number of human pathogens like *M. tuberculosis* [24], *Burkholderia pseudomalleii* [25], *Helicobacter pylori* [26] *Pseudomonas aeruginosa* [27], *Neisseria gonorrhea* [28] and *Salmonella typhi* [29]. In case of targeting genes or proteins for drug targets through subtractive and comparative microbial genome analysis, the main theme is to find such targets (gene/protein) that are essential for the pathogen and possess no homology counterpart in the host [30]. Along with subtractive genomics approach, reverse vaccinology strategy optimizes the prediction and development of novel drug and vaccine targets, especially for microorganisms that are difficult to grow in the laboratory, such as intracellular bacteria including *L. pneumophila*. Conventional vaccines may take more than 15 years to develop relying on adequate antigen expression from in vitro culture models including undesirable consequences while epitope-based vaccine prediction could be a faster approach targeting immunogenic protein through entire bacterial or viral proteome [31-33]. Reverse vaccinology approach usually identified a protein vaccine candidate using defined features, such as protein subcellular localization, topology, adhesion/antigenicity probability, epitopes, and its binding to the major histocompatibility complex (MHC) class I and II molecule [34]. Reverse vaccinology approach has been proven to prioritize and design vaccine targets against multiple pathogens [35-39].

In this study, we took in-depth subtractive genomics and reverse vaccinology approach to identify novel therapeutic drug and vaccine target in *L. pneumophila* subsp. *pneumophila (strain Philadelphia 1 / ATCC 33152 / DSM 7513).* We particularly considered the key essential or survival proteins of the pathogen which are non-homologous to the host as well as host microbiota and screened for outer membrane proteins (OMP) and B/T-cell epitopes. At the same time, we performed the human microbiome non-homologous analysis. The highest scoring OMP’s and epitopes will facilitate future in vitro and in vivo tests for the production of drugs and suitable vaccine against intracellular *L. pneumophila* infection.

## Materials and methods

The genome-wide proteome exploration of *L. pneumophila* was employed for identifying novel drug and vaccine targets through subtractive genomics and reverse vaccinology approach. The overall workflow has been illustrated in Fig. 1 and Fig. 2.

**Fig. 1.** Proteome exploration of *Legionella pneumophila* to identify novel drug targets.

**Fig. 2.** Flow chart summarizing the protocols for the prediction of epitope based vaccine candidate by *in silico* reverse vaccinology technique.

### Proteome retrieval and Finding paralogous sequences

The entire proteome of *L. pneumophila* subsp. *pneumophila (strain Philadelphia 1 / ATCC 33152 / DSM 7513)* (UniProt ID: UP000000609), was retrieved from UniProt(http://www.uniprot.org/proteomes/) containing total 2930 proteins [40] which were subjected to CD-Hit analysis (http://weizhongli-lab.org/cdhit_suite/cgi-bin/index.cgi?cmd=cd-hit) [41]. Sequence identity cutoff score was set at 0.6 to exclude redundant sequences of more than 60% identity through this method. Only non-paralogous protein sequences excluding the redundant sequences (paralogous sequences) were passed through further study.

### Identification of human (host) non-homologous proteins

The goal of the non-homology analysis of *L. pneumophila* subsp. *pneumophila* is to identify pathogen specific-proteins which are non-homologous to the human. BLASTp analysis was carried out on non-paralogous proteins against the reference proteome of *Homo sapiens* using threshold expectation value (E value) 10^−3^ through Ensembl (https://uswest.ensembl.org/Multi/Tools/Blast?db=core) [42]. Proteins that showed significant hits were filtered out meaning that they had similarities with the human genome leaving just the nonhomologous sequences. These non-homologous sequences were screened. The purpose of this step is to avoid any functional resemblance with human proteomes and to reduce any unwanted cross-reactivity of the drugs thereby avoiding the binding of drugs to the active site of homologous human proteins [43].

### Identification of essential nonhomologous proteins

Nonhomologous proteins were subjected to Database of Essential Genes (DEG) [44]. DEG contains 53,885 essential genes and 786 essential non-coding sequences. BLASTp of previously selected nonhomologous proteins was carried out by selecting all the organisms present in DEG 15.2 server and using threshold expectation value 10^−10^, a minimum bit-score cut-off of 100 as parameters, but the hits with e-value ≤10^−100^, identity ≥ 25%, and same annotated function of the query was selected as essential proteins. These essential proteins were employed for further study. Proteins of essential genes in a bacterium shape a minimal genome, containing a set of functional modules that play key roles in the evolving area, synthetic biology [44].

### Metabolic pathway analysis and family prediction of hypothetical proteins

Kyoto Encyclopedia of Genes and Genomes [KEGG] includes full metabolic pathways in living organisms [45]. Using three letter KEGG organism code hsa and lpn of human and *Legionella pneumophila subsp. Pneumophila*respectively, all the metabolic pathways present in the host (*H. sapiens*) and the pathogen were collected separately. A manual comparison was made to identify the metabolic pathways that were only present in the pathogen, as unique to *Legionella pneumophila* subsp. *pneumophila* using KEGG PATHWAY Database [46] whereas the remaining pathways were grouped as common. The predicted human nonhomologous, essential proteins of *Legionella pneumophila subsp.pneumophila* were screened by BLASTp through KAAS [47] server at KEGG for the identification of potential drug and vaccine targets. KAAS (KEGG Automatic Annotation Server) generates functional annotation of genes by BLAST comparisons against the manually compiled KEGG GENES database and metabolic proteins are listed by assignments of KO (KEGG Orthology). It automatically generates KEGG pathways which indicate certain metabolic proteins. For the next phase of subcellular localization study, proteins involved in these particular metabolic pathways of the pathogen and proteins allocated to KO but not involved in specific pathways, were identified, except those proteins involved in common human and pathogen pathways. SVMProt is a server that uses Support Vector Machine to identify a protein sequence from its primary sequence into a specific class that contains all major groups of enzymes, transporters, receptors, pathways, RNA-binding and DNA-binding proteins. TheSVMProt server[48] (http:/bidd2.nus.edu.sg/cgi-bin/svmprot/svmprot.cgi) was used to determine the functional classes of the pathogen’s specific metabolic proteins in all uncharacterized, hypothesized proteins.

### Prediction of subcellular localization

For the reasons of gram-negativity, *Legionella pneumophila subsp.* proteins can be identified in five viable subcellular locations, such as cytoplasm, inner membrane, periplasm, outer membrane and extracellular. Cytoplasmic proteins can be used as targets for medications while surface membrane proteins can be used as targets for both the medication and vaccine[49]. PSORTb v3.0.2 [50] (http:/www.psort.org/psortb/index.html), CELLO v.2.5 [51] (http:/cello.life.nctu.edu.tw/), ngLOC (http:/genome.unmc.edu/ngLOC/index.html) was used to predict the location of selected non-homologous essential pathogen proteins. The best score for a location was counted, generated by these servers. Final location of these proteins was taken through three steps: 1) if all three servers predicted the same location for a protein, that location was selected as final result, 2) if any two of these three servers predict the same location for a protein, that location was selected as final result, 3) if they predicted three different locations for a protein then PSLpred server [52] (http://crdd.osdd.net/raghava/pslpred/) was used and the predicted location for that protein done through PSLpred server was matched with previous results predicted by PSORTb v3.0.2, CELLO v.2.5, and ngLOC servers.

### Druggability Screening and antigenicity analysis

DrugBank Database 5.1.0 contains entries including 2556 approved small molecule drugs, 962 licensed pharmaceutical drugs, 112 nutraceuticals, and more than 5125 experimental drugs. In addition, 5117 non-redundant protein sequences (i.e. drug target / enzyme / carrier / transporter) are linked to these drug entries. A ‘druggable’ target must have the potential to bind to drugs and drug-like molecules with high affinity. Shortlisted cytoplasmic and membrane proteins were screened through the database of DrugBank 5.1.0[53] using default parameters. Presence of drug targets from the shortlisted cytoplasmic and membrane proteins in DrugBank database indicating same biological functions can act as an evidence for their druggable properties and were grouped as current therapeutic targets while their absence indicated the novelty of those drug targets, hence were classified as ‘novel therapeutic targets’. Total 45 proteins that showed no similarity after passed through DrugBank database were listed as novel drug targets and vaccine candidates. VaxiJen v2.0 [54] was used for predicting the protective antigens and subunit vaccines. Only membrane protein sequences of novel drug targets of *Legionella pneumophila* subsp. *Pneumophila* were subjected to VaxiJen v2.0 selecting threshold value 0.4. Proteins that showed antigenicity prediction >0.4 in VaxiJen v2.0 were transferred for next step as potential vaccine candidates.

### ‘Anti-target’ analysis of the essential, nonhomologous and novel drug targets

For humans, the human Ether-a-go-go Related Gene (hERG), constitutive androstane receptor (CAR), pregene X receptor (PXR), and P-glycoproteins (P-gp) are anti-targets or alternative drug targets for host protein candidates. Some receptive membrane are aelso classified as ‘anti-target’. they are adrenergicα1a, dopaminergic D2, muscarinic M1, and serotonergic 5-HT2C. A total of 210 human ‘anti-targets’ have been identified in the literature (Supplementary file S1)[11] and the corresponding sequences of these proteins have been obtained from the NCBI Protein database. BLASTp was carried out for all nonhomologous, essential ‘novel drug target’ proteins against these ‘anti-targets’ setting an E-value threshold <0.005, query length >30%, identity <30% as parameters. Nonhomologous proteins showing <30% identity against these ‘anti-target’ proteins were transferred to next step.

### Human microbiome non-homology analysis

The correlation between gut flora and human is not merely commensal but rather is symbiotic, mutualistic relationship [59], and different beneficial functions by human microbiome were also been reported [60,61].Unintentional blocking or accidental inhibition of proteins present in thismicroflora may lead to adverse effects in the host [62]. Screening of nonhomologous, essential proteins selected as vaccine candidates and novel drug targets were subjected to BLASTp through NCBI protein blast server (https://blast.ncbi.nlm.nih.gov/Blast.cgi) using an E-value ‘cutoff score 1’ against the dataset present in Human Microbiome Project server (https://www.hmpdacc.org/hmp/) “43021 (BioProject)” [63] was mentioned in the Entrez Query field while selecting Non-redundant protein sequences (nr) as the database. The Human Microbiome Project (HMP) collected microbial strains from a disease-free, stable, 239-person adult population, which included 18 body habitats in five areas (oral, skin, nasal, gut, and urogenital), creating 5026 microbial species compositions[64]. Proteins showing < 48% similarity were identified as novel therapeutic targets and vaccine candidates and moved to the next stage. The results of this screening study ensured the chance to prevent unintended inhibition and unconscious blockage of human microflora proteins.

### Protein-Protein Interaction studies

The protein-protein interaction studies (PPIs) of selected shortlisted proteins were predicted using STRING v10.5 [65]. The database identifies both physical (direct) and functional (indirect) interactions. PPIs with high confidence score (≥70%) were considered to avoid false positive results. Inhibition of essential proteins can hamper other proteins to perform correctly. Therefore, analysisof PPIs of the shortlisted proteins with others of the same strain can help to identify the best therapeutic targets. Proteins showing close interactions with at least three others were selected for further studies. Only outer membrane protein with antigenicity score >0.6 was transferred to the next stage for reverse vaccinologyafter homology modeling and validation step and the rests were listed as probable novel drug targets.

### Homology modeling, structure optimization and validation

Since no exact PDB structure was available for the selected proteins, a BLASTp was carried out in NCBI for each protein where Protein Data Bank Proteins (pdb) was selected as the Database. The templates were chosen for homology modeling considering sequence identity ≥85 and query cover ≥30. EasyModeller 4.0 software was used to generate the best 3D structure of each selected therapeutic proteins[66]. The best models were subjected to ModRefiner for energy minimization and structure refinement [67]. Root mean square deviation (RMSD) values were also calculated. Evaluation of the optimized, refined structures was carried through online servers, PROCHECK [68] and ERRAT [69] present in Structural Analysis and Verification Server (SAVES) (http://servicesn.mbi.ucla.edu/SAVES/) to get the stereochemical quality of the model.

### Antigenic protein selection and T-Cell epitope prediction

Among the membrane associated proteins, the final vaccine target was selected according to protein-protein interaction analysis and antigenicity. ProtParam [70] tool showed different physico-chemical parameters of the proteins. The IEDB MHC-I and MHC-II prediction tools were used to predict MHC-I and MHC-II binding peptides, respectively. [71].

### Transmembrane topology, antigenicity, population coverage and allergenicity assessment of T-Cell epitopes

The TMHMM server predicted transmembrane helices in protein[72], while the VaxiJen v2.0 server was used to test antigenicity of the epitopes[54]. Population coverage has been evaluated for each human epitope using an IEDB analysis tool[70]. Four independent servers i.e. AllerTOP[73], AllergenFP[74], PA3P[75], and Allermatch[76] have been used to estimate epitope allergicity for potential vaccine design.

### Toxicity and conservancy analysis

The server ToxinPred (http:/crdd.osdd.net/raghava/toxinpred/) estimated the relative toxicity of top T-cell epitopes. Conservancy of the epitope has been shown to determine the degree of distribution in the homologous protein set of corresponding epitopes using the IEDB epitope conservancy analysis tool (http:/tools.iedb.org/conservancy/).

### Identification of B-Cell epitopes and cluster analysis

Six separate IEDB algorithms, that is. The Kolaskar and Tongaonkar antigenicity scale[77], the Karplus and Schulz versatility prediction[78], the Chou and Fasman beta turn prediction[79], the Emini surface accessibility prediction[80], the Parker hydrophilicity prediction[81] and the Bepipred linear epitope prediction tool[82] were used to identify the most active B cell epitopes in the target protein. Additionally, an analysis tool for the IEDB epitope cluster was used to identify overlapping peptides among the predicted epitopes[83]. Between top epitopes of CTL, HTL and BCL, larger cassettes containing multiple epitopes were found.

### Final vaccine construction

The final vaccine was formulated using the top epitopes (T-cell and B-cell), the adjuvants and the corresponding linkers. Three specific adjuvants, i.e. beta-defensin, ribosomal protein L7/L12 and M. Tuberculosis HABA protein (AGV15514.1) has been used to produce three separate vaccine molecules. Adjuvants interact with toll-like receptors (TLRs) and cause immune activation by polarizing CTL reactions[84]. Beta defensin adjuvant is TLR1, 2 and 4 agonist, whereas L7/L12 ribosomal protein and HBHA protein act only as TLR4 agonist. EAAAK, GGGS, GPGPG and KK linkers have been used to bind the adjuvant, CTL, HTL and B-cell epitopes, respectively. Literature reports also state that linkers allow efficient in vivo separation of individual epitopes[85,86]. To that the problem with highly polymorphic HLAs, the PADRE sequence was also inserted into vaccine constructs.

### Allergenicity, antigenicity, physico-chemical property and secondary structure analysis of vaccine constructs

AlgPred v.2.0 [87] sever was used to predict the non-allergic nature of the constructed vaccines. In order to suggest the superior vaccine candidate, VaxiJen v2.0 server[54] was used to evaluate its antigenicity. ProtParam[70,88] has shown molecular weight, instability index, approximate half-life, isoelectric pH, GRAVY values, hydropathicity, aliphatic index and other physico-chemical properties of the constructs. The secondary structure of the vaccine protein was determined by PSIPRED v3.3[89] and NetTurnP v1.0 [90,91].

### Vaccine tertiary structure prediction, refinement, validation and disulfide engineering

The RaptorX server generated the tertiary structure of the constructs based on the degree of similarity between target protein and accessible template structure in PDB database[92,93]. Refinement was conducted using ModRefiner [94] followed by FG-MD refinement server [95] to improve the accuracy of the predicted 3D modeled structure. Ramachandran plot of the refined structure was assessed by RAMPAGE [96]. Disulfide bonds enhance the geometric conformation of proteins and provide significant stability. DbD2 was used to build these bonds for the vaccine designs [97]. The value of chi3 selected for the residue screening was -87 to +97, while the energy value considered was less than 2.5.

### Protein-protein docking and molecular dynamics simulation

Inflammations caused by bacterial antigen are involved with TLR-2 immune receptors present over the immune cells [98-100]. The 3D structure of different MHC molecules and human TLR-2 receptors has been obtained from the database of RCSB proteins. Molecular docking method using ClusPro[101], hdoc[102,103] and PatchDock server[104] determined the binding affinity of designed vaccines with specific HLAs and TLR-2 immune receptors. FireDock server refined the complexes generated via PatchDock server [104]. The vaccine-receptor complex’s stability was determined by contrasting the critical protein dynamics to their normal modes[105,106]. Essential dynamics is a versatile tool and an inexpensive solution to costly atomistic simulation [106,107]. iMODS server explained the collective motion of the proteins by analyzing the normal modes (NMA) in internal coordinates [108,109]. The server was used as it assess much faster than other molecular dynamics simulations tools [110,111]. The direction and extent of the immanent motions of the complex was analyzed in terms of B-factors, deformability, covariance and eigenvalue [112].

### Codon adaptation and in silico cloning

For greater expression of the vaccine protein in *E. coli*, a codon adaptation technique was used. The procedure was conducted by the JCAT server[113], while preventing the termination of Rho’s independent transcription, the ribosome-binding prokaryote site, and the cleavage of several other restriction enzymes (i.e. HindIII and BamHI). The optimized sequence of vaccine protein V1 was reversed and then conjugated with HindIII and BamHI restriction site at the N-terminal and C-terminal sites respectively. SnapGene restriction cloning module was used to insert the adapted sequence between HindIII (173) and BamHI (198) of pET28a(+) vector [114].

## Results

All the methodologies used in the proteome exploration of *Legionella pneumophila* including the number of proteins screened in each step had been summarized in Table 1.

**Table 1:**
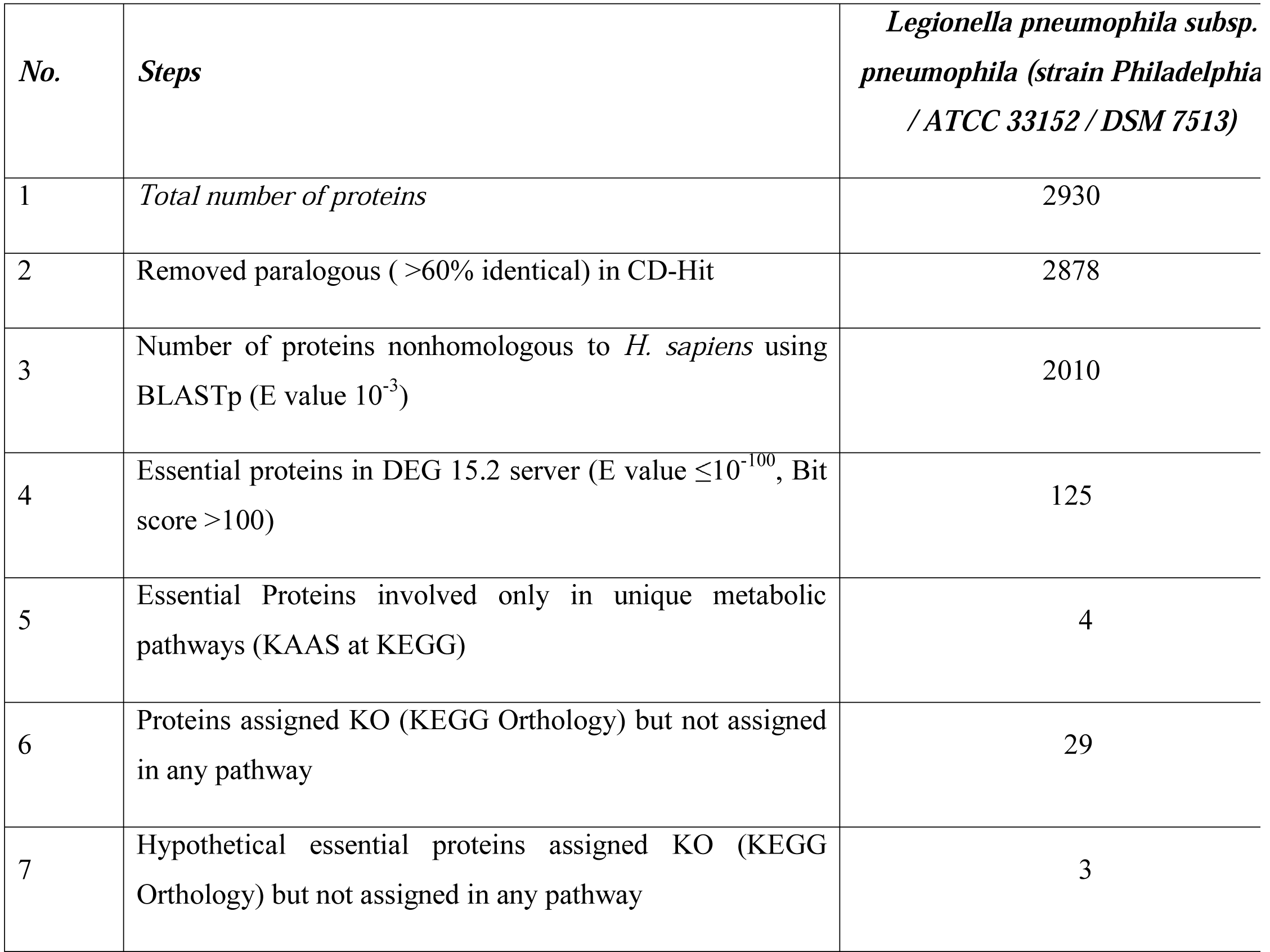
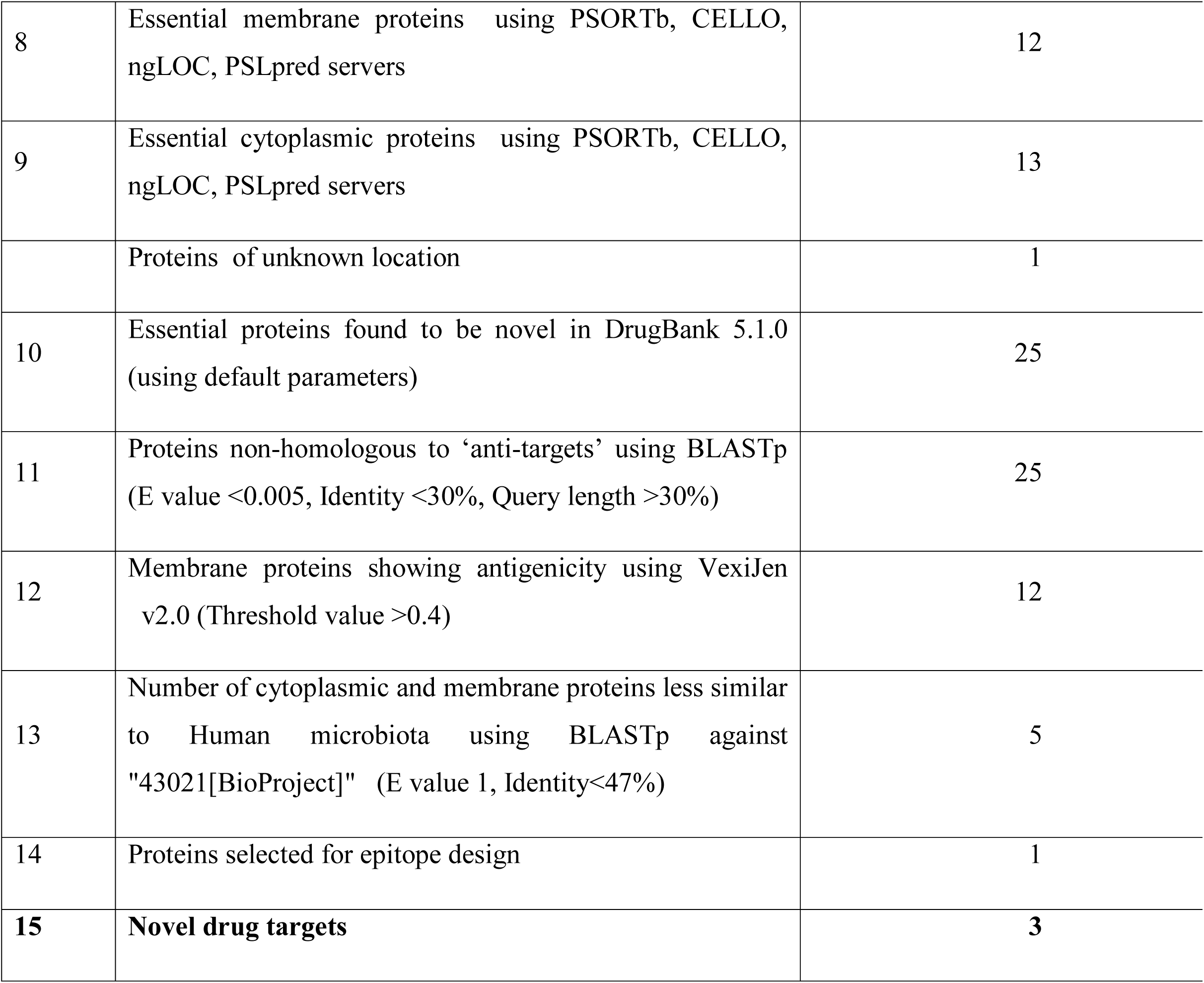
Subtractive genomic analysis scheme towards the identification of novel therapeutic targets.

| *No.* | *Steps* | *Legionella pneumophila subsp. pneumophila (strain Philadelphia / ATCC 33152 / DSM 7513)* |
| --- | --- | --- |
| 1 | *Total number of proteins* | 2930 |
| 2 | Removed paralogous ( >60% identical) in CD-Hit | 2878 |
| 3 | Number of proteins nonhomologous to *H. sapiens* using BLASTp (E value $10^{-3}$) | 2010 |
| 4 | Essential proteins in DEG 15.2 server (E value $\leq 10^{-100}$, Bit score >100) | 125 |
| 5 | Essential Proteins involved only in unique metabolic pathways (KAAS at KEGG) | 4 |
| 6 | Proteins assigned KO (KEGG Orthology) but not assigned in any pathway | 29 |
| 7 | Hypothetical essential proteins assigned KO (KEGG Orthology) but not assigned in any pathway | 3 |
| 8 | Essential membrane proteins using PSORTb, CELLO, ngLOC, PSLpred servers | 12 |
| 9 | Essential cytoplasmic proteins using PSORTb, CELLO, ngLOC, PSLpred servers | 13 |
|  | Proteins of unknown location | 1 |
| 10 | Essential proteins found to be novel in DrugBank 5.1.0 (using default parameters) | 25 |
| 11 | Proteins non-homologous to ‘anti-targets’ using BLASTp (E value <0.005, Identity <30%, Query length >30%) | 25 |
| 12 | Membrane proteins showing antigenicity using VexiJen v2.0 (Threshold value >0.4) | 12 |
| 13 | Number of cytoplasmic and membrane proteins less similar to Human microbiota using BLASTp against "43021[BioProject]" (E value 1, Identity<47%) | 5 |
| 14 | Proteins selected for epitope design | 1 |
| **15** | **Novel drug targets** | **3** |

### Proteome retrieval and Finding paralogous sequences

The whole proteome retrieved from UniProt (UniProtID: UP000000609) containing 2930 proteins were analyzed in CD-Hit tool using threshold value 0.6 to eliminate paralogous and duplicate sequences showing >60% sequence identity. Paralogous protein sequences were removed leaving total 2878 non-paralogous protein sequences.

### Identification of human non-homologous proteins

Non-paralogous protein sequences were screened by BLASTp through Ensembl genome browser 92 868 proteins was found showing similarity with the human genome which were excluded leaving total 2010 non-homologous proteins.

### Identification of essential proteins

Among 2010 nonhomologous proteins, only 125 protein sequences showed similarity with the essential proteins enlisted in DEG server from 48 bacterial strains. These 125 proteins were listed as essential proteins and are considered as responsible for the survival of *Legionella pneumophila.*

### Metabolic pathway analysis and family prediction of hypothetical proteins

Though 120 pathways of target *Legionella pneumophila* were identified from KEGG server, only 37 pathways were detected as unique for pathogen (Supplementary file S2). The result from KAAS server at KEGG revealed that 122 proteins among 125 had assigned KO identifier. From 122, only 4 proteins were involved in unique metabolic pathways (Table 2) and 29 proteins involved neither in unique nor common pathways (Supplementary Table 1). Only 3 hypothetical proteins were found among these 29 KO assigned proteins for which functions remains unknown. Functional family for these hypothetical proteins were also analyzed (Supplementary Table 2). All of these 33 proteins were allowed for further investigation. Remaining 89 proteins were excluded since these were found to be involved in both pathogen and human metabolic pathways.

**Table 2:** Proteins involved in pathogen specific pathways.

| *S. no.* | *KO assignment* | *Accession no.* | *Protein name* | *Pathway name* |
| --- | --- | --- | --- | --- |
| 1 | *K18138* | tr\|Q5ZXL1 | Multidrug resistance protein | beta-Lactam resistance; Cationic antimicrobial peptide (CAMP) resistance |
| 2 | *K01928* | tr\|Q5ZX16 | UDP-N-acetylmuramoyl-L-alanyl-D-glutamate--2,6-diaminopimelate ligase | Lysine biosynthesis; Peptidoglycan biosynthesis |
| 3 | *K03587* | tr\|Q5ZX17 | Peptidoglycan D,D-transpeptidaseFtsI | Peptidoglycan biosynthesis; beta-Lactam resistance |
| 4 | *K05515* | tr\|Q5ZVR5 | Peptidoglycan D,D-transpeptidaseMrdA | Peptidoglycan biosynthesis; beta-Lactam resistance |

### Prediction of subcellular localization

The shortlisted 33 nonhomologous essential proteins involved in unique pathways of the pathogen were screened through PSORTb, CELLO and ngLOC servers. A total 16 proteins were identified as cytoplasmic proteins and 16 proteins were identified as membrane proteins which were further screened for druggability and antigenicity analysis (Supplementary file S3). But, the study failed to predict the subcellular localization of ‘tr|Q5ZYU2’ since these four servers generated different predictions.

### Druggability Screening and antigenicity analysis

Only 7 proteins showed similarity with the available drug targets (Table 3) at DrugBank5.1.0 database. The remaining 25 showed no hitand thesewere considered as novel drug targets of *L. pneumophila* which were comprised of 13 cytoplasmic and 12 membrane proteins. Antigenicity analysis of membrane protein revealed that only 1 protein was non antigenic whereas the rest 11 showed antigenicity score more than 0.4. at VaxiJen server (Table 4). Furthermore, all of the proteins screened after druggability and antigenicity analysis were employed for human ‘anti-targets’ and microbiome non-homology study.

**Table 3:** Identified druggable targets with drug names.

| *Sl. No* | *Accession No.* | *Protein name* | *Drugbank Id* | *Drug name* | *Bit score* | *Query length* |
| --- | --- | --- | --- | --- | --- | --- |
| 1 | tr\|Q5ZS52 | RNA polymerase sigma factor RpoH | DB08874 | <u>Fidaxomicin</u> | 103.605 | 318 |
| 2 | tr\|Q5ZT04 | RNA polymerase sigma factor RpoD | DB08226,<br>DB08266 | <u>Myxopyronin B,</u><br><u>Methyl [(1E,5R)-5-{3-[(2E,4E)-2,5-dimethyl-2,4-octadienoyl]-2,4-dioxo-3,4-dihydro-2H-pyran-6-yl}hexylidene]carbamate</u> | 302.368 | 248 |
| 3 | tr\|Q5ZUP0 | UDP-N-acetylmuramate--L-alanyl-gamma-D-glutamyl-meso-2,6-diaminoheptandioate ligase | DB01673,<br>DB03909,<br>DB04395 | <u>Uridine-5'-Diphosphate-N-Acetylmuramoyl-L-Alanine,</u><br><u>Adenosine-5'-[Beta, Gamma-Methylene]Triphosphate,</u><br><u>Phosphoaminophosphonic Acid-Adenylate Ester</u> | 145.591 | 398 |
| 4 | tr\|Q5ZXL1 | Multidrug resistance protein | DB03825,<br>DB04209,<br>DB03619 | <u>Rhodamine 6G</u>,<br><u>Dequalinium</u>,<br><u>Deoxycholic Acid</u> | 427.557 | 1020 |
| 5 | tr\|Q5ZX17 | Peptidoglycan D,D-transpeptidaseFtsI | DB00303,<br>DB00671 | <u>Ertapenem</u>,<br><u>Cefixime</u> | 122.094 | 632 |
| 6 | tr\|Q5ZVR5 | Peptidoglycan D,D-transpeptidaseMrdA | DB01598,<br>DB01329,<br>DB01327,<br>DB01163,<br>DB01328,<br>DB01413,<br>DB01415,<br>DB00948,<br>DB00438,<br>DB00303,<br>DB06211 | <u>Imipenem</u>,<br><u>Cefoperazone</u>,<br><u>Cefazolin</u>,<br><u>Amdinocillin</u>,<br><u>Cefonicid</u>,<br><u>Cefepime</u>,<br><u>Ceftibuten</u>,<br><u>Mezlocillin</u>,<br><u>Ceftazidime</u>,<br><u>Ertapenem</u>,<br><u>Doripenem</u> | 486.493 | 612 |
| 7 | tr\|Q5ZZI4 | Amino acid permease | DB00123,<br>DB00125,<br>DB00129 | <u>L-Lysine</u>,<br><u>L-Arginine</u>,<br><u>Ornithine</u> | 155.606 | 424 |

**Table 4:** List of probable antigenic proteins for vaccine targets.

| *S. no.* | *Accession no.* | *Vaxijen v2.0 score* | *Subcellular localization* |
| --- | --- | --- | --- |
| 1 | tr\|Q5ZS84 | 0.4733 | Inner Membrane |
| 2 | tr\|Q5ZUM7 | 0.4869 | Inner Membrane |
| 3 | tr\|Q5ZT44 | 0.5906 | Inner Membrane |
| 4 | tr\|Q5ZVR6 | 0.6058 | Inner Membrane |
| 5 | sp\|Q5ZV69 | 0.6714 | Outer membrane |
| 6 | sp\|Q5ZW98 | 0.6393 | Inner Membrane |
| 7 | sp\|Q5ZSY2 | 0.4632 | Inner Membrane |
| 8 | sp\|Q5ZTN5 | 0.5461 | Inner Membrane |
| 9 | sp\|Q5ZRL2 | 0.4190 | Inner Membrane |
| 10 | tr\|Q5ZY66 | 0.4935 | Outer membrane |
| 11 | tr\|Q5ZX71 | 0.5807 | Inner Membrane |

### ‘Anti-target’ and Human microbiome non-homology analysis

To avoid severe cross-reaction and toxic effects in human, identification of nonhomologous proteins to human essential proteins (referred as ‘anti-targets’) is a crucial step. Study revealed that all selected 25novel therapeutic proteins showed no similarity with 210 human ‘anti-targets’. In addition, BLASTp against all microbial strains presented in Human Microbiome Project (HMP) server carried out through NCBI blast server revealed that only 5 proteins out of 25 showed similarity ≤47% (Table 5). Targeting these proteins will be suitable since they are neither involved in common pathways of host-pathogen nor homologous to any human ‘anti-targets’.

**Table 5:** Human microflora non-homology analysis and subcellular localization.

| S. no. | Accession no. | Microbiome similarity | Subcellular localization |
| --- | --- | --- | --- |
| 1 | tr\|Q5ZX16 | <46 | Cytoplasmic |
| 2 | sp\|Q5ZUD8 | <45 | Cytoplasmic |
| 3 | tr\|Q5ZS84 | <47 | Inner Membrane |
| 4 | sp\|Q5ZV69 | <47 | Outer membrane |
| **5** | **sp\|Q5ZW98** | **<44** | **Inner Membrane** |

### Protein-Protein Interaction studies

PPIs in STRING v10.5 revealed that Cytoplasmic protein ‘UDP-N-acetylmuramoyl-L-alanyl-D-glutamate--2,6-diaminopimelate ligase (murE)’ showed interactions with proteins involved in cell wall biosynthesis pathways (Fig. 3A) whereas ‘Trigger factor (tig)’ showed interactions with proteins which were mainly involved in binding with different rRNA and tRNA molecules (Fig. 3B). Moreover, inner membrane protein ‘Probable lipid II flippaseMurJ (mviN)’ exhibited interactions with proteins which were connected to peptidoglycan biosynthesis pathway and transportation of lipid-linked peptidoglycan precursors (Fig. 3C), where ‘Biotin synthase (bioB)’ is an essential protein for the survival of pathogen. The name of essential drug targets listed in Table 6. Another inner membrane protein ‘Probable potassium transport system protein kup 1 (kup1)’ presented single interaction (Fig. 3D) and as a consequence it was excluded from our analysis. Moreover, the remaining outer membrane protein ‘TolB (tolB)’ showed close interactions with different other proteins (Fig. 3E) which were mainly found in peptidoglycan synthesis and lipoproteins translocation process. ‘Protein TolB’ also exhibited less similarity (<47%) with human microflora proteins and had antigenicity score >0.6, that is why, it had been considered as potential vaccine candidate against *Legionella pneumophila*. So, further study was employed for ‘in silico vaccine design’ focusing ‘Protein TolB’ by reverse vaccinology approach. Details of PPIs was provided in Supplementary file S4.

**Fig. 3.** Investigation of Protein-Protein Interactions throughSTRING v10.5 server, **A:** UDP-N-acetylmuramoyl-L-alanyl-D-glutamate--2,6-diaminopimelate ligase (murE), **B:** Trigger factor (tig), **C:** Lipid II flippaseMurJ (mviN), **D:** Potassium transport system protein kup 1 (kup1), **E:**Protein TolB (tolB).

**Table 6:** List of identified novel drug targets against *Legionella pneumophila*.

| *S. no.* | *Accession no.* | *Protein names* |
| --- | --- | --- |
| 1 | tr\|Q5ZX16 | UDP-N-acetylmuramoyl-L-alanyl-D-glutamate--2,6-diaminopimelate ligase |
| 2 | sp\|Q5ZUD8 | Trigger factor |
| 3 | tr\|Q5ZS84 | Probable lipid II flippaseMurJ |

### Homology modeling, structure optimization and validation

The selected single vaccine candidate Q5ZV69 was modeled to determine 3D structure through EasyModeller 4.0 (Supplementary Fig. 1A). Complete results of template selection for homology modeling are provided in Supplementary Table 3. The best structure generated for each target was optimized by ModRefiner server. Refined model of Q5ZV69 had RMSD 0.1444 and TM score 0.9783 to initial model. After refinement, the PDB structure of each protein was evaluated through PROCHECK and ERRAT. Psi-Phi pains in Ramachandran Plot revealed that protein Q5ZV69 had 91.9% residues in most favored regions, 7.8% residues in additional allowed region and 0.3% in generously allowed region (Supplementary Fig. 1B). According to ERRAT, the overall quality factor of this protein was 76.1084 (Supplementary Fig. 2). An ERRAT score of 50 is normally acceptable and the value explains the statistics of non-bonded interaction molecules. This indicates that the modeled structure is reliable and stable.

### Antigenic protein selection and T-Cell epitope prediction

From the top membrane proteins, LEGPH Protein TolB (Accession ID: Q5ZV69) was selected based on protein-protein interaction analysis with total antigenicity score of 0.6740 for vaccine design against *Legionella*. ProtParam tool was used for analyzing the physicochemical properties of the vaccine proteins. Results of which are shown in Supplementary Table 4. In addition, various immunogenic epitopes from LEGPH Protein TolB were identified to be potential T cell epitopes that can bind to wide variety of different HLA-A and HLA-B cells with greater binding affinity. Epitopes that bind to the maximum number of HLA cells were selected.

### Transmembrane topology, antigenicity, population coverage & allergenicity assessment of T-T-cell epitopes

Best LEGPH Protein TolB epitopes were ranked based on their topology of the transmembrane and antigenicity score (Table 7). All of the indicated alleles were identified as optimal binders of the proposed epitopes and used to assess population coverage (Fig. 4). Based on studies utilizing four separate allergenicity prediction systems, epitopes classified as non-allergenic to humans have been retained in the predicted epitope list (Table 7).

**Fig. 4.** Population coverage analysis of LEGPH protein TolB.

**Table 7:**
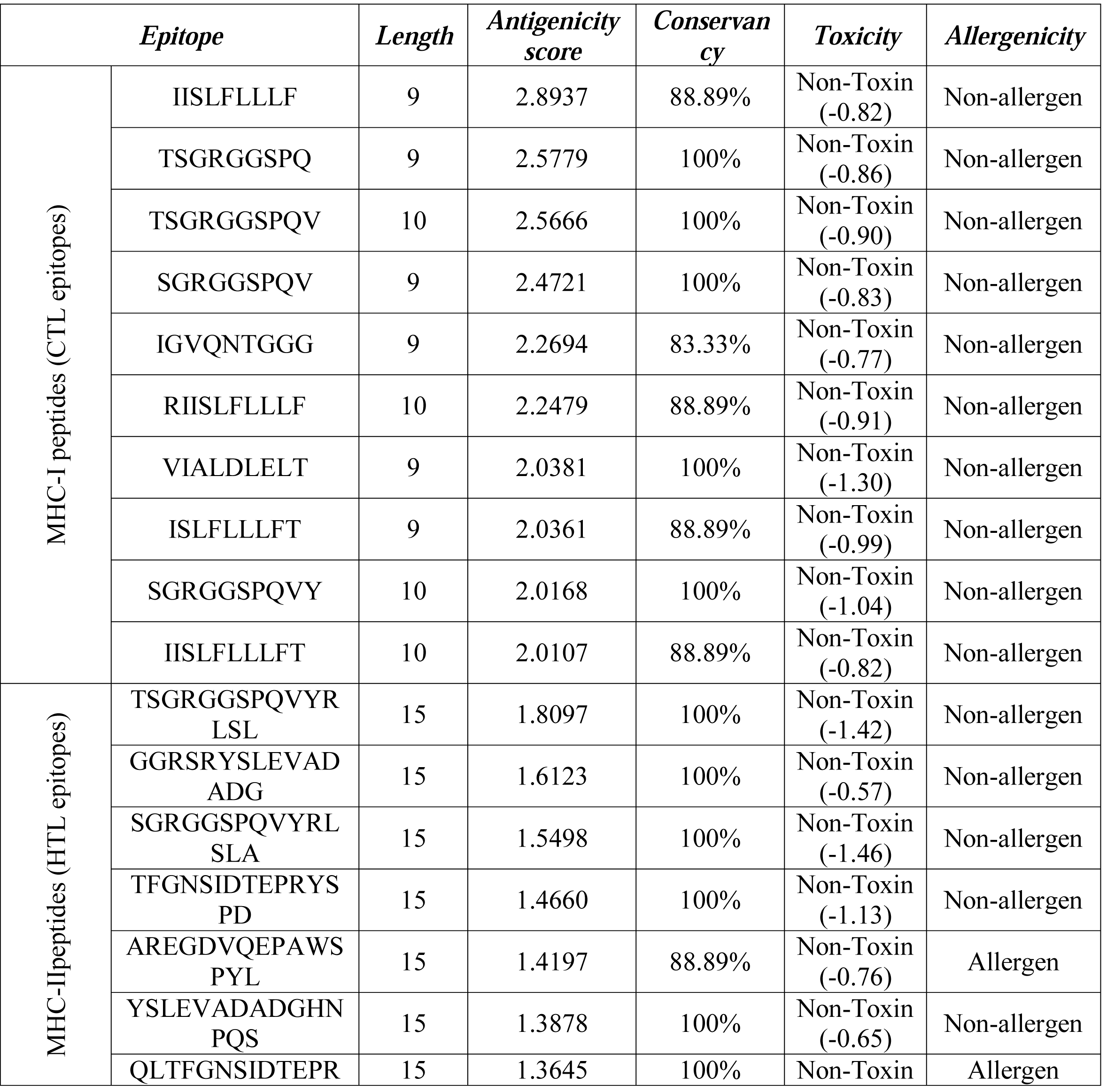
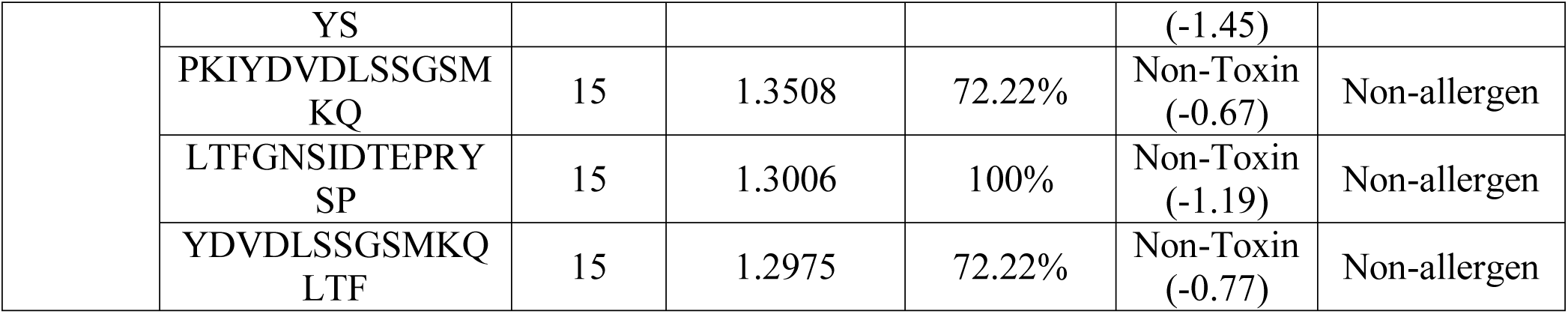
Predicted T-cell epitopes (MHC-I peptides and MHC-II peptides) of LEGPH protein TolB.

### Toxicity and conservancy analysis of T-Cell epitope

ToxinPred server predicted the relative toxicity of the top epitopes (Table7). Again, putative epitopes generated from LEGPH Protein TolB were found to be highly conserved (up to 100%) within specific pathogenic strains. Top epitopes were used to design the final vaccine constructs to allow a wide spectrum of immunity (Table 7).

### B-Cell epitope identification and cluster analysis

B-cell epitopes of the selected protein was generated using a total six algorithms (Table 8). For envelope glycoprotein, peptide sequences from 66-77 and 404-417 were identified as potential B cell epitopes that could stimulate preferred immune reaction (Supplementary Fig. 3A). Regions from 206-212 and 277-287 residues were more accessible (Supplementary Figure 3B), while residues from 194-200 and 294-300 amino acids were considered as possible regions of Beta-turn (Supplementary Fig. 3C). On the basis of Karplus and Schulz flexibility prediction, the region of 160-166 and 308-314 residues were most flexible (Supplementary Fig. 3D). Amino acids in 5-23 and 243-252 regions were highly antigenic (Supplementary Figure 3E), while residues in the 174-180 and 283-289 regions were mostly hydrophilic (Supplementary Fig. 3F). Before the final constructs were designed, peptides containing overlapping epitopes between both the top T-cells and B-cells were identified (Supplementary Table 5).

**Table 8:**
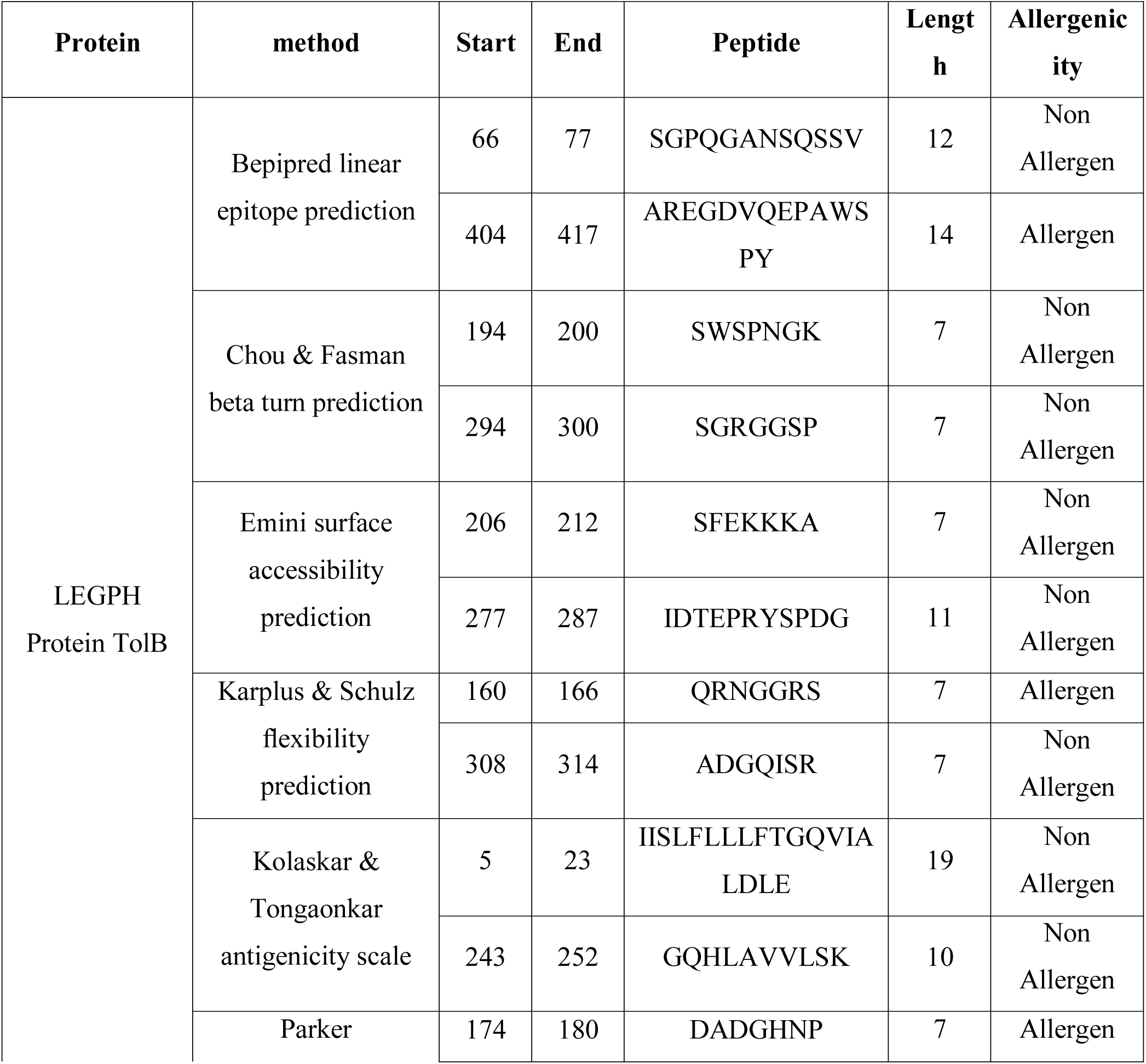
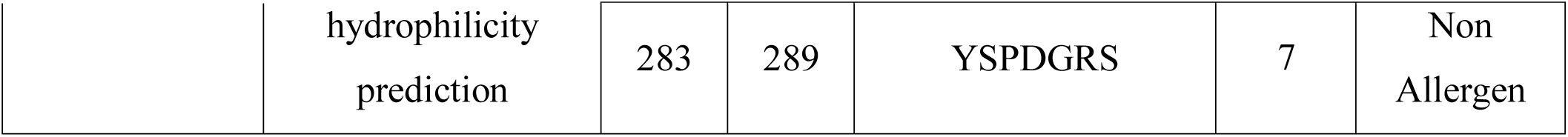
Allergenicity pattern of the predicted B-cell epitopes generated from LEGPH protein TolB.

### Final vaccine construction

Both Clusters and singletons (contains single epitope) were used to construct the final vaccine molecules. A total of 3 constructs with residues with 476 (V1), 561 (V2), and 590 (V3) amino acids were designed (Table 9) and used for analysis.

**Table 9:**
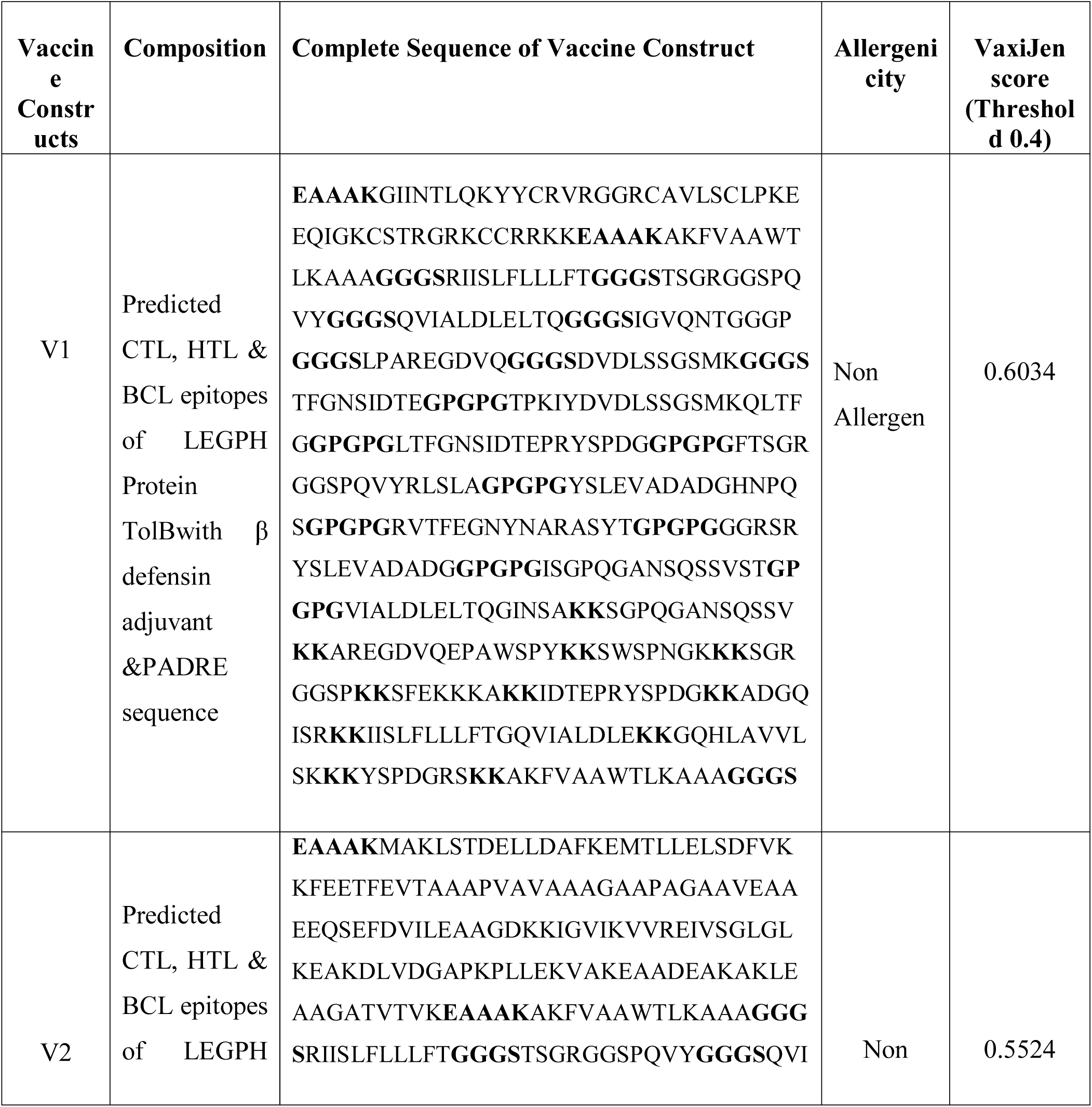
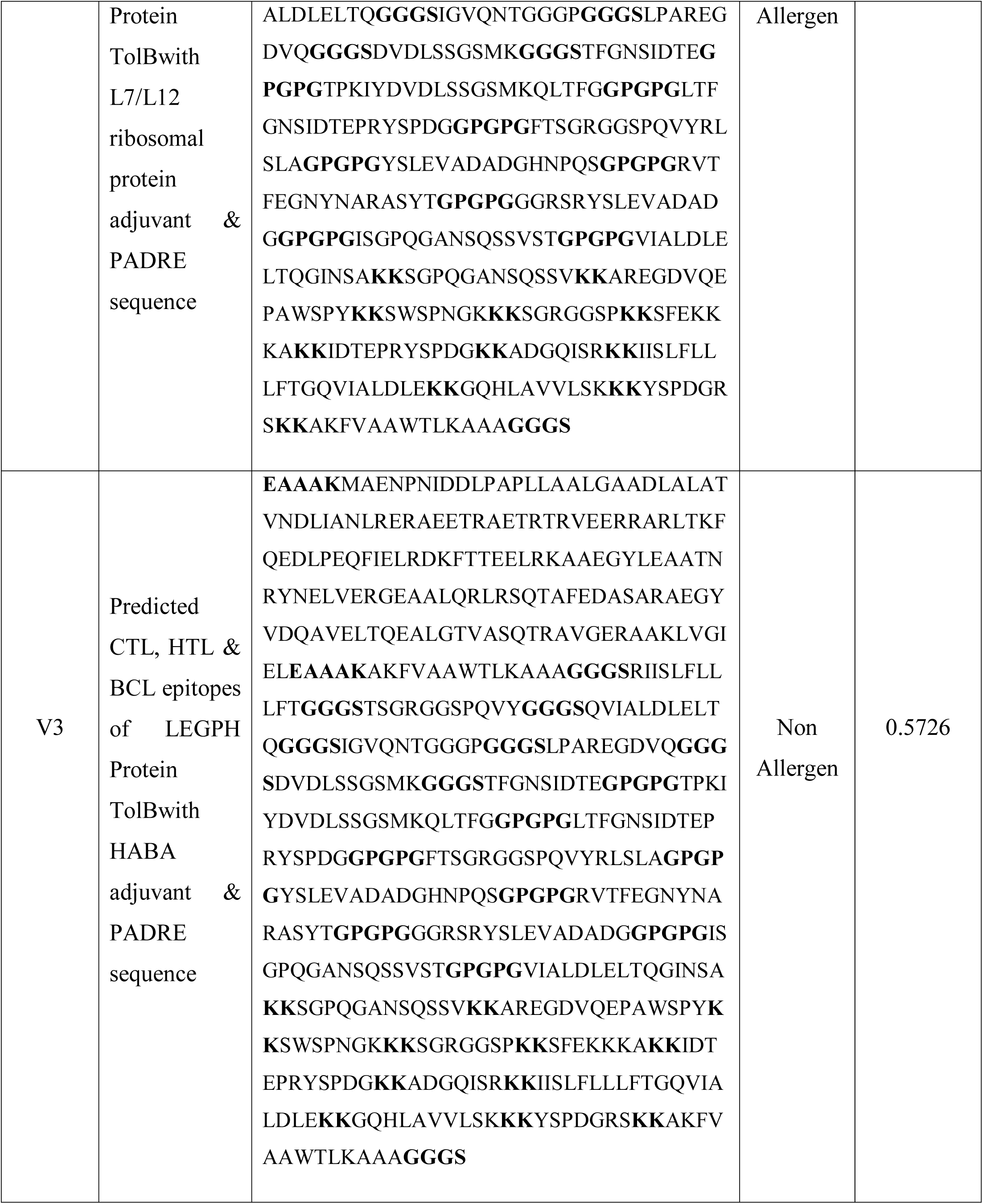
Allergenicity prediction and antigenicity analysis of designed vaccine constructs.

**Table 10:** Docking score of vaccine construct V1 with different HLA alleles including HLA-DRB1*03:01 (1A6A), (HLA-DRB5*01:01 (1H15), HLA-DRB3*01:01 (2Q6W), HLA-DRB1*04:01 (2SEB), HLA-DRB1*01:01 (2FSE), and HLA-DRB3*02:02 (3C5J)

| Vaccine Construct | PDB ID of the HLA Alleles | Global Energy | Hydrogen Bond Energy | ACE | Score | Area |
| --- | --- | --- | --- | --- | --- | --- |
| V1 | 1A6A | -12.98 | -4.93 | 7.18 | 17648 | 3028.30 |
|  | 1H15 | -30.47 | -5.55 | 5.82 | 16608 | 2414.80 |
|  | 2Q6W | -10.99 | -2.58 | 7.65 | 17294 | 2662.20 |
|  | 2SEB | -27.99 | -8.91 | 0.83 | 18036 | 2334.50 |
|  | 2FSE | -1.43 | -0.46 | 0.05 | 19568 | 2748.70 |
|  | 3C5J | -9.43 | -3.63 | 15.90 | 16902 | 2337.90 |

### Allergenicity, antigenicity, physicochemical properties and secondary structure analysis of vaccine constructs

Results revealed construct V1’s superiority due to greater antigenicity score (0.603) and non-allergenic behavior (Table 9). The final construct of the vaccine was characterized by the physical and chemical properties. Construct V1’s molecular weight was 48.34kDa, while theoretical pI was measured at 9.78 which indicated that the protein should have a net negative charge over the pI and vice versa. The half-life of the vaccine was expected to be more than 10 h in *E. coli* in vivo. The estimated rate of extinction and an aliphatic index were 41370 and 61.74 respectively. The protein’s computed GRAVY value was -0.545 while the index of instability (33.51) classified the protein as stable. Constructed V1 was, on the opposite, characterized by 6.30% alpha helix, 24.78% sheet and 68.92% coil structure (Supplementary Fig. 4).

### Vaccine tertiary structure prediction, refinement, validation and disulfide engineering

2ymuA from the PDB database was detected as the most suitable V1 template and a single-domain 3D model was created through the Raptor X server (Fig. 5A). All 476 residues were modelled by the server, while only 8% residues were in the disordered region. The 3D model’s P value was 5.30e-11 which ensured better quality modeling of the proposed vaccine. 81.4 %, 11.8 % and 6.8 % residues were respectively in the favored, allowed and outlier regions prior to refinement. However, 88.4% residues were in the favored region after refining. Ramachandran Plot showed residues of 8.4 % and 3.2 % respectively in the allowed and outlier regions (Fig. 5B). Homology modelling of construct V2 and V3 was also performed as shown in Supplementary Fig. 5. For construct V1, 71 amino acid pairs were found to have the capacity to shape disulfide bond. However, just 3 pairs i.e., based on the chi3 and B-factor values ASN 202-GLY 226, SER 211-GLY 217 and ASN 309-LEU 317 met disulfide bond forming requirements (Fig. 6).

**Fig. 5.** Tertiary structure prediction and validation of vaccine protein V1, **A:** Cartoon format **B:** Validation of the 3D structure of vaccine protein V1 by Ramachandran plot analysis.

**Fig. 6.** Disulfide engineering of vaccine protein V1; **A:** Initial form, **B:** Mutant form.

### Protein-protein docking and molecular dynamics simulation

Docking study revealed that construct V1 bound in the groove of different HLAs with minimal binding energy which was biologically significant. The predicted binding energy for vaccine V1-TLR 2 complex was -1232.8 and -275.55 KJule/mol by ClusPro and hdoc server respectively. FireDock output refinement of PatchDock server showed the lowest global energy of -15.64 for solution 2. The lowest binding energy was a measure of highest binding affinity between TLR-2 and vaccine construct. The stability of proteins and their mobility at large scale were revealed by normal mode analysis (NMA). Deformabilty of each residue was negligible and represented by hinges in the chain (Fig. 8A). The B-factor value was equivalent to RMS inferred via NMA (Fig. 8B). The eigenvalue found for the complex was 3.5007e-06 (Fig. 8C). The eigenvalue and variance associated to each normal mode were inversely related (Fig. 8D). The covariance matrix indicated the coupling between pairs i.e. correlated (red), anti-correlated (blue) or uncorrelated (white) motions (Fig. 8E). The elastic network model detected the pairs of residues which were connected via springs (Fig. 8F).

**Fig. 7.** Docked complex of vaccine construct V1 with human TLR8; **A:** Cartoon format and **B:** Ball structure.

**Fig. 8.** Molecular dynamics simulation of vaccine protein V1-TLR2 complex; stability of the protein-protein complex was investigated through mobility **(A)**, eigenvalue **(B)**, B-factor **(C)**, deformability **(D)**, covariance **(E)** and elastic network **(F)** analysis.

### Codon adaptation for in silico cloning and expression in E. coli

The optimized codon adaptation index (CAI) was found 0.95 and GC content of that swquence was 54.01%. This results showed better expression in *E. coli* K12 The optimized codons were introduced into the vector pET28a(+) along with the restriction sites HindIII and BamHI. A clone of 6346 base pair was created in which red color indicated the desired sequence of 1434 bp in pET28a(+) vector sequence (Fig. 9).

**Fig. 9.** Restriction digestion **(A)** and *in silico* cloning **(B)** of the gene sequence of final vaccine construct V1 into pET28a(+) expression vector. Target sequence was inserted between HindIII (173) and BamHI (198).

## Discussion

Among all the pathogenic agents responsible for pneumonia and respiratory tract infection, *Legonella pneumophila* is one of the main causative agent showing high levels of resistance against a large number of antibiotics. Though clinical studies of antibiotic treatment against Legionnaires’ disease have been reported but the number of such cases are not quite enough [115]. Hence, Hence, treatment recommendations are limited due to lack of evidence [116]. However, Rahimi and Vesalreported that *L. pneumophila* strains harbored highest levels of resistance againstciprofloxacin, erythromycin, clarithromycin, moxifloxacin andazithromycin [117]. Therefore, the prime goal of this study was to screen out and determine novel drug targets and vaccine candidates for *Legionella pneumophila* through subtractive genomics and reverese vaccinology approach.

Computational predictions help to identify essential proteins for the survival of the pathogen, non-homologous to host proteins, as well as no involvement in metabolic pathways of host leading to choose the proteins only present and involve in pathogen specific metabolic pathways. Complete reference proteome of *L. pneumophila* (2,930 proteins) was searched and downloaded from the NCBI protein database. Proteins involved in several common cellular systems emerged as homologous with same functions between human and bacteria in course of evolution [118,119] which were removed in this study based on their identity with human proteins. Essential proteins are most promising for new antibacterial drug targets since most antibiotics are designed to bind essential gene products [120] and can be considered as pathogen-specific drug targets [121]. The study revealed 125 unique essential proteins of *L. pneumophila* which can be considered as suitable drug targets. Localization is an important aspect of any possible drug target as cellular functions of proteins are often localized in specific compartments of cell, hence studying subcellular localization will be helpful to understand disease mechanisms as well as developing new vaccine candidates and drug targets. Although, both membrane and cytoplasmic proteins could serve the purpose as therapeutic targets [122], membrane proteins habe been mostly reported for vaccine candidates [123]. Hence, in this study membrane proteins were used for vaccine construction whereas cytoplasmic proteins were proposed as suitable drug targets. Again, usage of antibiotics has already reduced their efficiency due to gene mutation and thereby rapid emergence of resistant bacteria [124]. [124]. The antibiotic and antimicrobial resistance crises are observed due to the misuse as well as overuse of these drugs, and a scarcity of developing new drugs by the pharmaceutical industry [125,126]. Broad-spectrum drugs medicated for a pathogen or group of the pathogen may cause mutational changes as well as enhance the transfer of gene to other pathogens which can show resistance to drugs leading towards the emergence of resistant bacteria. To avoid these crises, selection of novel drug targets and vaccine candidates is a must; therefore shortlisted 32 proteins were further screened through DrugBank5.1.0 database with default parameters.

Antigenicity score for selected 12 novel drug targeted membrane proteins revealed that only 1 protein was non antigenic whereas the rest 11 showed antigenicity score >0.4. These 11 proteins can be used to design B cell epiptopes and T cell epitopes (specific for CTL and HTL) in future to lessen the disease caused by *Legionella pneumophila*. A number of drugs from different categories such as non-nacrotic analgesics, non-steroidal anti-inflammatory drugs, antidepressants as well as vasodilators were withdrawn during 1960-1999 due to various health-relatedcross-reactivity causing hepatic, cardiovascular, dermatologist and hematologic toxicities as well as carcinogenic effects [127]. For example, bromfenac [128], ebrotidine [129], trovafloxacin [130] were withdrawn from pharmaceutical markets worldwide since they showed hepatotoxic effects. Therefore, specific recognition must be maintained by an ideal drug target to the drug-treated against it, furthermore, this drug target needs to be significantly different from the host proteins. To avoid severe cross-reaction and adverse effects in human, identification of nonhomologous proteins to human ‘anti-targets’ (also refferend as essential protein) was a crucial step considered in this study. Targeting Human microbiome non-homology proteins will be suitable since they are neither involved in common pathways of host-pathogen nor homologous to any human ‘anti-targets’.Furthermore drug or vaccine designed and administered for these novel targets will be less harmful to other commensal microbial strains dwelling in different body sites of a healthy human. The study revealed that most of the cytoplasmic proteins are involved in cell wall biosynthesis pathways and essential for cell division. However, targeting three proteins i.e. UDP-N-acetylmuramoyl-L-alanyl-D-glutamate--2,6-diaminopimelate ligase, Trigger factor and Probable lipid II flippaseMurJ also influence the interacting proteins can lead to development of novel therapeutics in The predicted single vaccine candidate Q5ZV69 of this study was analyzed to develop a potential and highly immunogenic vaccine candidate against *Legionella pneumophila*. Using a variety of bioinfomatics tools, various antigenic epitopes were generated which were extensively investigated for immunogenicity, toxicity profile, allergenicity pattern and conservancy analysis The PADRE sequence reportedly decreased the polymorphism of HLAs in the different population [131,132]. In previous in vivo research, the linkers improved immunogenicity of the vaccines [133,134]. Therefore, in the present study, both of these critical considerations were taken into account in developing the final vaccine model Moreover, the results of docking analysis revealed the binding affinity of promiscuous epitopes with different HLA alleles. In addition, interaction between construct V1 and TLR-2 were checked to demonstrate the efficacy of used adjuvants. The vaccine-receptor complex showed minimum deformability as well at molecular level.

## Conclusion

By using the subtractive genomics and reverse vaccinology approach, we are trying to develop novel therapeutics against *Legionella pneumophila* and may help to reduce the rate of mortality as well as morbidity caused by this pathogen. We particularly considered the key that are the must for survival of the pathogen, non-homologous to the human as well as human microbiota and screened out B/T-cell epitopes of outer membrane protein (OMP). The highest scoring OMP’s and epitopes will facilitate future wet lab based experiments for the development of drugs and suitable vaccine candidates against intracellular *L. pneumophila* infection. After intensive analysis, only 3 proteins were identified and proposed as novel therapeutic targets against *L. pneumophila*. Only outer membrane protein TolB was identified as potential vaccine candidate with a better antigenicity score. However, we also suggest further *in vitro* and *in vivo* laboratory trials to validate our prediction.

## Supplementary Figures

**Supplementary Fig. 1.** Tertiary structure prediction of the vaccine target through EasyModeller 4.0 **(A)** and Ramachandran Plot analysis **(B)**.

**Supplementary Fig. 2.** Quality factor analysis of modeled 3D structure of LEGPH protein TolB by ERRAT.

**Supplementary Fig. 3.** Prediction of B cell linear epitope and intrinsic properties for LEGPH protein Tol B using different scales (**A:** Bepipred, **B:** Surface accessibility, **C:**Emini surface, **D:** Flexibility, **E:** Antigenicity, **F:** Hydrophilicity). (For each graph: x-axis and y-axis represent the position and score; residues that fall above the threshold value are shown in yellow color; the highest peak in yellow color identifies most favored position).

**Supplementary Fig. 4.** Secondary structure prediction of designed vaccine V1 using PESIPRED server

**Supplementary Fig. 5.** 3D modelled structure of vaccine protein V2 **(A)** and V3 **(B)** generated via RaptorX server.

## Supplementary Files

**Supplementary File 1:** A total 210 ‘anti-targets’ of human

**Supplementary File 2**: Unique pathways of *Legionella pneumophila* retrieved from KEGG server.

**Supplementary File 3:** Subcellular localization of nonhomologous essential proteins involved in unique pathways of *Legionella pneumophila*

**Supplementary File 4:** Details of Protein-Protein Interactions through STRING v10.

**Supplementary Table 1:**
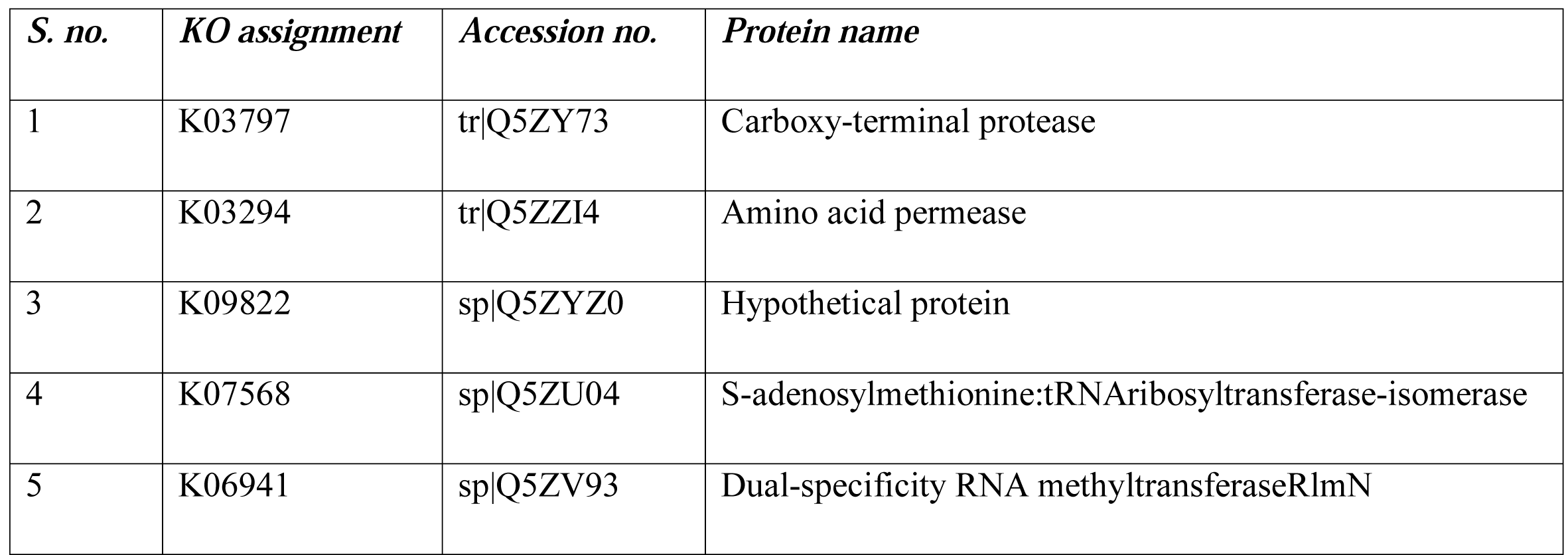
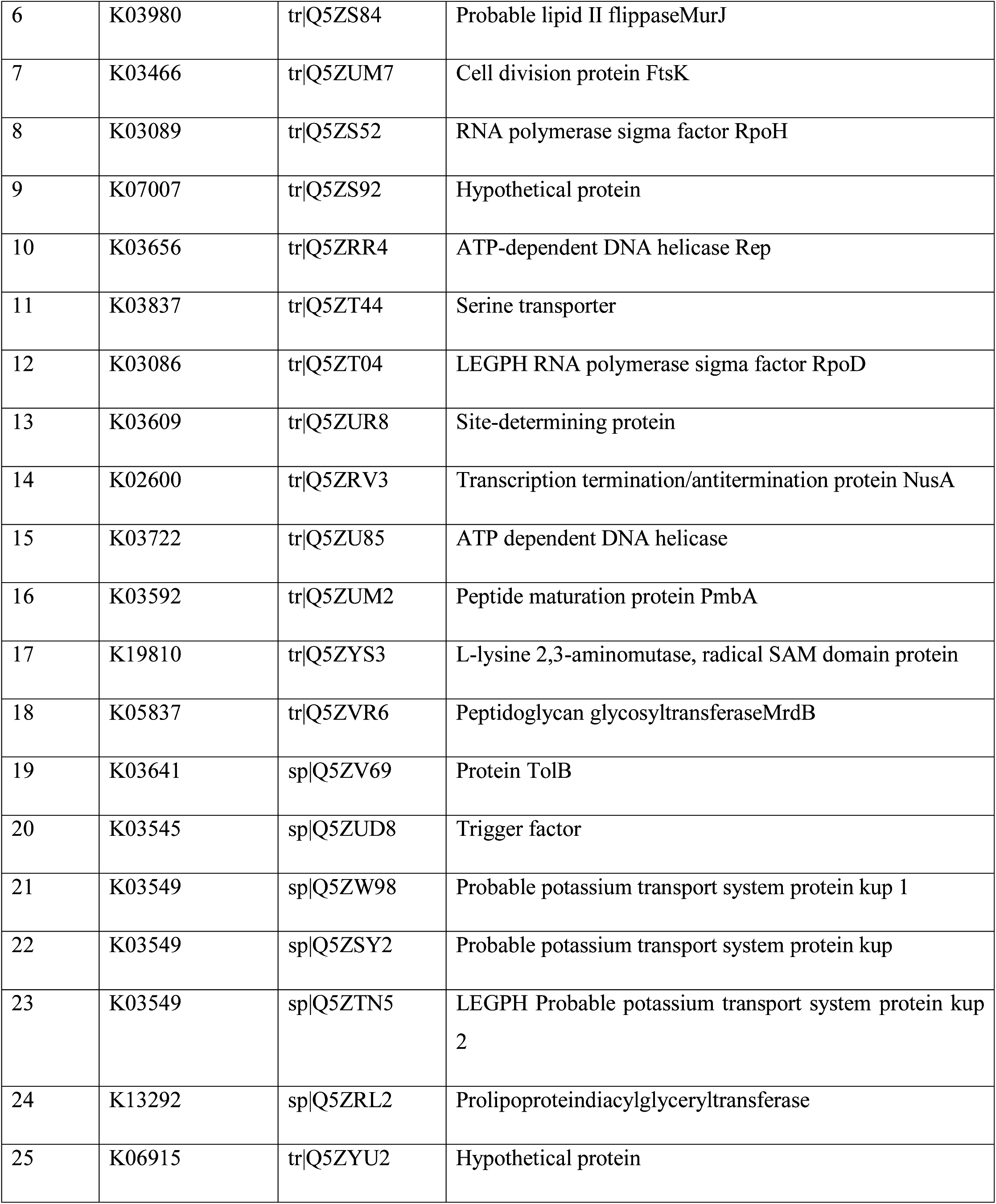
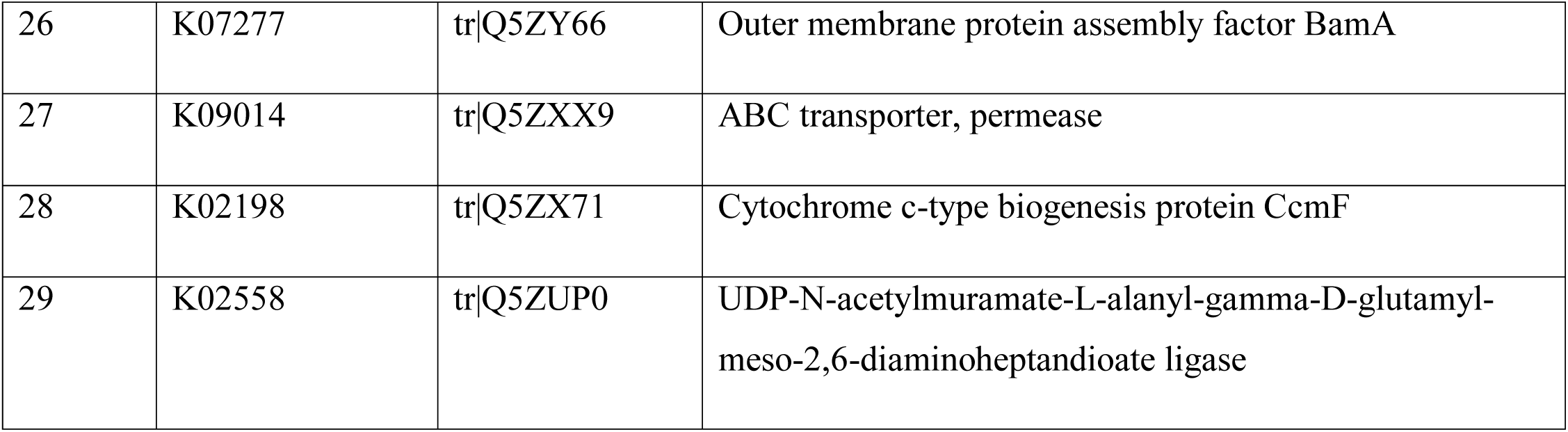
KO assigned (KEGG orthlogy) proteins not involved in any metabolic pathway.

**Supplementary Table 2:** Functional family classification of the hypothetical proteins by SVMProt server.

| *S. no.* | *Accession no.* | *KO assignment* | *Family name* | *Probability (%)* | *GO category* |
| --- | --- | --- | --- | --- | --- |
| 1 | sp\|Q5ZYZ0 | K09822 | Iron-binding | 98.4 | GO:0005506 |
| 2 | tr\|Q5ZS92 | K07007 | Iron-binding | 98.9 | GO:0005506 |
| 3 | tr\|Q5ZYU2 | K06915 | All DNA-binding | 85.4 | GO:0003677 |

**Supplementary Table 3:** Templates used for homology modeling by EasyModeller 4.0.

| Protein accession no. | PDB ID | Identity (%) | Query cover |
| --- | --- | --- | --- |
|  | 4R40 | 41 | 95 |
| sp\|Q5ZV69 | 3IAX | 40 | 99 |
|  | 1C5K | 40 | 99 |

**Supplementary Table 4:** ProtParam analysis of LEGPH protein TolB.

| *Viral Proteins* | *Accession ID* | *Molecular Weight* | *Instability Index* | *Aliphatic Index* | *Theoretical pI* | *No. of Amino acids* | *Extinction Co-Efficient* | *Gravy* |
| --- | --- | --- | --- | --- | --- | --- | --- | --- |
| TolB_LEGPH | Q5ZV69 | 45387.06 | 38.09 | 89.81 | 8.67 | 419 | 34380 | -0.204 |

**Supplementary Table 5:**
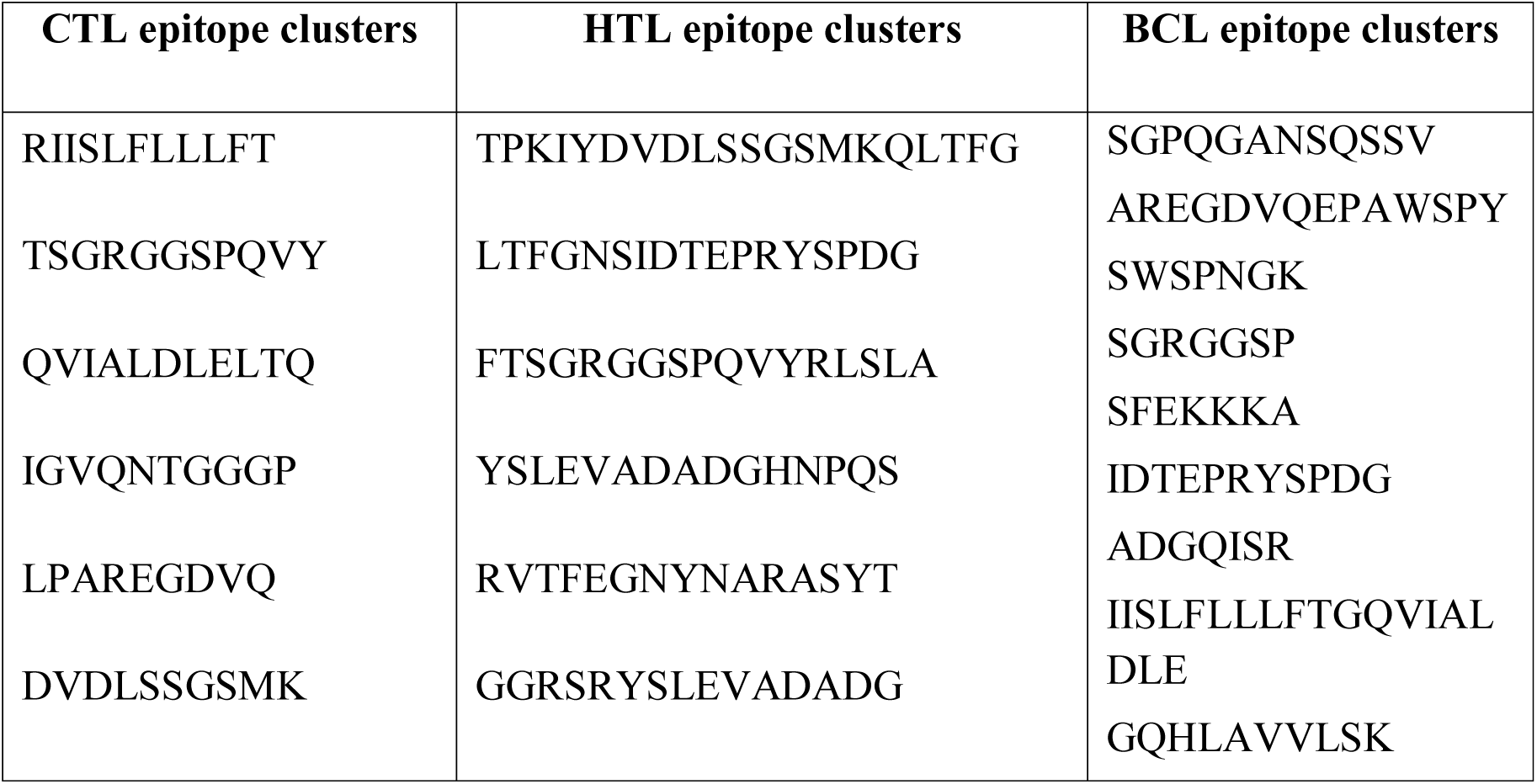
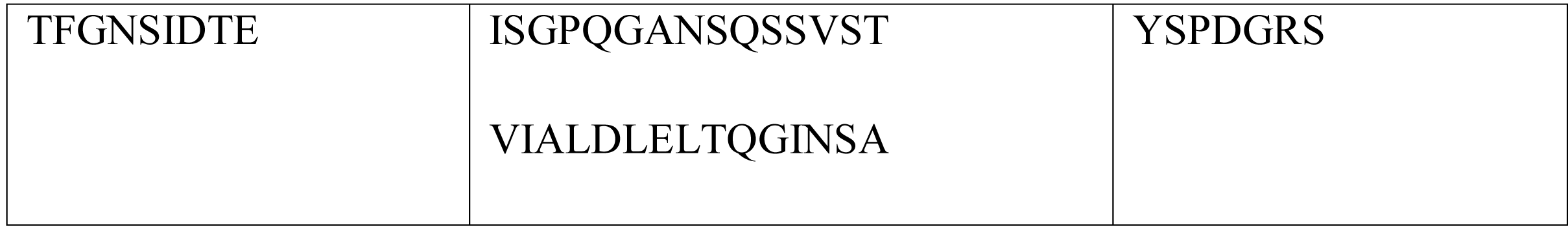
Identified epitope clusters among top CTL, HTL and BCL epitopes.

